# *In vitro* evolution of uropathogenic *Escherichia coli* to fosfomycin resistance in a 3D cultured human bladder microtissue model

**DOI:** 10.64898/2026.07.22.740011

**Authors:** Beth James, M J Wilde, Macsen T Fryer, Benjamin O Murray, Daniel J Whiley, Charlotte Cornbill, Jennifer L Rohn, Alasdair T M Hubbard

**Author notes:** corresponding author: Alasdair Hubbard.

## Abstract

*In vitro* studies of antimicrobial resistance (AMR) using laboratory growth media produce important, fundamental information. However, their inability to more closely replicate the *in vivo* environment limits the translational potential of this work. Here, we used a 3D cultured microtissue model which reflects the human bladder microenvironment to select for resistance to fosfomycin in two uropathogenic strains of *Escherichia coli*, UTI-34 and UTI-59. To assess the clinical relevance of the mutations produced, we screened the observed mutations in the fosfomycin-selected variants against a curated dataset of 14,163 *E. coli* genomes isolated from urine. The four independent fosfomycin-selected variants of UTI-34 contained diverse mutations, while the mutations in the five independent fosfomycin-selected variants of UTI-59 were more constrained. All variants contained mutations in *glpT*, *uhpT*, *uhpA* and *uhpC*, which are commonly linked to fosfomycin-resistance in clinical isolates of *E. coli*. Screening of the mutations against the 14,163 *E. coli* genomes from urine confirmed that four of these mutations were found as exact matches in the dataset, while other mutation types were confirmed at a regional and gene level. These mutations did not result in any collateral susceptibility or resistance to other antibiotics recommended for the treatment of urinary tract infections. The use of a human 3D microtissue model, which closely replicates the urothelial microenvironment to study AMR during urinary tract infection treatment, could improve the clinical relevance of *in vitro* AMR studies. This has the potential to provide a better understanding of how AMR is acquired and expressed, and inform new strategies to combat AMR.

## Introduction

The acquisition and evolution of antimicrobial resistance (AMR) are widely studied *in vitro* using laboratory growth media[1–4]. Laboratory media often fail to recapitulate the infection environment, and even host-mimicking media does not accurately match the infection-like environment[5]. Therefore, due to the divergent conditions between laboratory media and the *in vivo* setting[6–8], there is a growing need to study AMR acquisition and evolution in more physiologically relevant conditions. One strategy is to use longitudinal collections of isolates with or without matched clinical prescribing data to map AMR acquisition and evolution[3, 9–11]. However, these datasets are limited, difficult to obtain and expensive. In addition, isolates have often been collected from a prior study, accessed retrospectively and are non-optimised for the current study[9–11]. *In vitro* models of specific infection sites exist, including *ex vivo* and 3D culture models, and are common in the study of host-pathogen interactions[12–18]. While these models have been used to study the efficacy of antimicrobials[12, 14], to date, there has been limited exploration of using them in the study of AMR acquisition and evolution.

One excellent candidate is the <u>3D</u> <u>U</u>rine-tolerant <u>H</u>uman <u>U</u>rothelial (3D-UHU) model, which closely mimics the human bladder urothelial microenvironment[17]. 3D-UHU is tolerant of 100% human urine for up to four weeks and stratifies to 7-8 cell layers with differentiation of basal, intermediate and umbrella cells layers, closely resembling the human urothelium[17]. The 3D-UHU model produces key biomarkers, including uroplakin proteins and cytokeratins, and demonstrates strong barrier function[17]. Importantly, the model secretes human cytokines and chemokines in response to infection with uropathogens, including IL-1b, IL-8 and IL-6[17]. The reproducibility of the model, access to the apical chamber and physiological relevance make the 3D-UHU model an attractive option to study AMR acquisition and evolution during the treatment of urinary tract infections (UTIs).

In the UK, nitrofurantoin and trimethoprim are recommended as the first-line antibiotics for lower UTIs in non-pregnant women 16 years old and over, with pivmecillinam and fosfomycin the recommended second-line antibiotics[19]. Resistance to nitrofurantoin in bacteriuria *Escherichia coli* is very low, at approximately 2%[20–22], but when resistance does develop it is primarily caused by inactivating mutations in *nfsA* and *nfsB*[23–25]. In contrast, trimethoprim resistance is found at approximately 30% in bacteriuria *E. coli*[21, 22] and driven by the acquisition of plasmid-borne dihydrofolate reductase[26]. Pivmecillinam resistance is primarily driven by β-lactamase overproduction, such as *ampC* or a penicillinase, or the acquisition of a β-lactamase, including TEM or OXA, with a resistance rate of 8.3% of bacteriuria *E. coli*[22]. Like nitrofurantoin, fosfomycin resistance in bacteriuria *E. coli* is relatively low at 3.6%[22]. While fosfomycin can be acquired horizontally via fosfomycin-inactivating enzymes, including FosA[27, 28], resistance by mutation is also common. Such mutations are commonly found in the glycerol-3-phosphate transporter (*glpT)*[29–32], glucose-6-phosphate transporter (*uhpT*)[29–32] and UhpT regulatory system (*uhpABC)*[30–32]. Mutations in adenylate cyclase (*cyaA*)[30, 33] and Enzyme I (*ptsI*)[30, 33] of the phosphoenolpyruvate-carbohydrate phosphotransferase system have also been identified, but their exact role in fosfomycin resistance is unclear.

Here, we sought to determine if it was possible to use physiologically relevant conditions to recapitulate clinically relevant resistance *in vitro* using an advanced 3D cultured microtissue model. To achieve this, we used the 3D-UHU model in an experimental evolution study to select for AMR to fosfomycin in *E. coli*. We selected fosfomycin due to resistance arising by mutation (unlike trimethoprim and pivmecillinam) in a diverse set of genes (in contrast to nitrofurantoin) while having a higher resistance rate. Next, using a bioinformatics approach, we explored if the observed mutations were representative of the diversity of mutations found in *E. coli* clinical isolates. We used genomes from the BakRep database due to the available metadata and the ability to easily search for specific criteria and uniformity in processing[34]. Finally, we assessed the collateral effects of three other antibiotics used for the treatment of UTIs in the UK, and the fitness of the ancestors and the fosfomycin-selected variants in Mueller Hinton (MH) broth.

## Materials and methods

### Bacterial strains and growth conditions

Two clinical isolates of *E. coli* isolated from urine, 19Y000034 (UTI-34) and 19Y000059 (UTI-59), were obtained from the Nottingham University Hospitals NHS Trust Pathogen Bank[4].

Unless otherwise stated, all bacteria were inoculated on MH agar (MilliporeSigma, USA) and incubated at 37°C for 18 hours. Starter cultures were made from a single bacterial colony in 10 ml MH broth (MilliporeSigma, USA) and incubated at 37°C, 150 rpm for 18 hours. Cultures were washed three times in PBS by centrifuging at 5,000 rpm for 5 minutes and the pellet was resuspended in 10 ml PBS.

Fosfomycin (Cambridge Biosciences, UK) and glucose-6-phosphate (Sigma, USA) were solubilised in deionised water at a stock concentration of 10 mg/ml and sterile-filtered through a 0.22 µM polyethersulfone filter unit (MilliporeSigma, USA).

### 3D-urine tolerant human urothelial model

The 3D-UHU was generated as previously described[18]. Briefly, 2D cultures of HBLAK (CELLnTEC, CH) were passaged up to passage 12, with passages 8-12 used for seeding the 3D model. Approximately 80-90% confluent cells were detached using Accutase (CELLnTEC, CH) and 4×10^5^ cells/ml were seeded onto 0.4 µM pore polycarbonate filter membranes in a 12-transwell plate (Corning, USA) in prewarmed CnT-Prime (CELLnTEC, CH). The transwell plate was incubated at 5% CO_2_ at 37°C for 48 hours and membranes were assessed for 100% confluency. When 100% confluent, the media in the apical and basal chambers were replaced with CnT-Prime-3D medium (CELLnTEC, CH), starting the 3D culture (day 0). The transwell plates were incubated under the same conditions for a further 24-48 hours, following which the media in the apical chamber was replaced with PHU and the basal chamber was replaced with fresh CnT-Prime-3D media. PHU was sourced commercially (BioIVT, USA) and pooled from 5 female and 5 male volunteers, sterilised with a 0.22 µM pore filter. The urine and CnT-Prime-3D media in the apical and basal chambers, respectively, were changed every 3-4 days until day 18, following which the models were used for experiments. One membrane insert per transwell plate was sacrificed as a control for confocal imaging to confirm the 3D stratification and differentiation of the 3D-UHU model.

### Confocal imaging

A membrane insert from each transwell was fixed with 4% methanol-free formaldehyde (Fisher Scientific, UK) in PBS for 15 minutes at room temperature, followed by two washes with PBS and one wash with HBSS. The 3D-UHU model was stained with 0.6µg/ml wheat germ agglutinate diluted in Hank’s balanced salt solution (HBSS; Sigma, US) and placed on a rocker at 50 rpm, covered in tin foil at room temperature, for an hour. The WGA was then aspirated and the membranes were washed with HBSS. The membranes were then excised with a scalpel and one placed in each well of a 24 well plate using tweezers. Then the cells were permeabilised with 0.2% Triton X-100 in PBS and placed on a rocker at 50 rpm, covered in tin foil at room temperature for 30 minutes. This was then aspirated and the cells washed with PBS. Cells were then stained with Phalloidin-555 (Abcam, UK) and 1 µg/ml 4′,6-diamidino-2-phenylindole (DAPI; Invitrogen, USA), both in PBS, and placed on a rocker at 50 rpm, covered in aluminium foil for 1 hour. The membrane insert was then washed 5 times in PBS and left at room temperature to dry for 1 minute. The membrane insert was mounted on a glass slide with mounting medium (Invitrogen, USA) and coverslip sealed with nail varnish. Samples were imaged using a Leica SP5 confocal microscope (Leica, DE) and analysed using Image J software (V1.54p).

### Antimicrobial susceptibility testing

A broth microdilution assay to assess the minimum inhibitory concentration (MIC) of fosfomycin against UTI-34 and UTI-59 was performed following EUCAST guidelines in MH broth, supplemented with 25 µg/ml glucose-6-phosphate. Additionally, broth microdilution assays were also performed in conditioned urine recovered from the apical chamber of the 3D-UHU model during differentiation (apical urine) from day 10 onwards, following the same protocol. MICs were confirmed visually. All assays were performed with three biological replicates.

Disc diffusion assays to assess the susceptibility of UTI-34, UTI-59 and the fosfomycin-selected variants to fosfomycin 200 µg, cephalexin 30 µg, nitrofurantoin 100 µg and trimethoprim 5 µg (all Oxoid, UK) were performed following EUCAST guidelines on MH agar. For fosfomycin, MH agar was supplemented with 25 µg/ml glucose-6-phosphate. Zones of inhibition were measured with a ruler. All assays were performed with three biological replicates.

### Selection for fosfomycin-selected variants

Five biological replicate cultures of UTI-34 or UTI-59 in PBS were diluted 1 in 10 into 270 µl of PHU, making a total volume of 300 µl, containing 0.25x MIC of fosfomycin and 25 µg/ml glucose-6-phosphate and added to the apical chamber of mature 3D-UHU models. The same concentration of fosfomycin (0.25x MIC) and 25 µg/ml glucose-6-phosphate was also added to the basal media. After 24 hours of incubation in 5% CO_2_ at 37°C, the Day 1 apical PHU was removed and replaced with 270 µl of fresh PHU containing 0.5x MIC of fosfomycin and 25 µg/ml glucose-6-phosphate. The Day 1 apical PHU was diluted 1 in 10 into the fresh PHU containing fosfomycin and glucose-6-phosphate to reseed with the culture (Day 2). The media from the basal chamber was removed and replaced with fresh media containing 0.5x MIC fosfomycin and 25 µg/ml glucose-6-phosphate. This was repeated for a third time, increasing the fosfomycin concentration to 1x MIC (Day 3). After a total of 72 hours of exposure to increasing concentrations of fosfomycin, 100 µl were plated onto MH agar containing 1x, 2x, 4x and 8x MIC of fosfomycin. Due to confluent growth, for UTI-34, fosfomycin-selected growth was re-streaked from 1x or 2x onto 4x and 8x to achieve single colonies. Single colonies from 8x MIC were re-streaked onto MH agar without antibiotics and a single colony was used to make a starter culture, which was then stored in MH broth with 10% glycerol at -80°C.

### Whole genome sequencing

DNA from all fosfomycin-selected variants was extracted using New England Biolabs Monarch Genomic DNA Extraction Kit (New England Biolabs, USA) following the manufacturer’s instructions, except DNA was eluted with molecular grade water (Sigma, USA). DNA concentration was quantified using a Qubit fluorometer and 1X dsDNA High Sensitivity (HS) assay kit (ThermoFisher Scientific, USA). Genome sequencing was provided by MicrobesNG (https://microbesng.com). All fosfomycin-selected variants were sequenced by using the Illumina platform with 2x 250 bp kits, and sequencing reads trimmed using Trimmomatic[35].

### Genome assembly and annotation

A hybrid assembly approach was used in which long-read sequencing data were first assembled into a consensus genome using Autocycler (version 0.1.0)[36], generating a high-quality draft assembly. Illumina paired-end reads were subsequently used to polish this long-read-derived consensus genome, as described below.

Trimmed Illumina paired-end reads were aligned to the Autocycler consensus assembly using minimap2 (version 2.28-r1209)[37] with the short-read preset (-x sr). The resulting alignments were used as input for polishing with Racon (version 1.5.0)[38], which was run with default parameters to generate an initial polished assembly.

The Racon-polished assembly was then indexed and used as a reference for short-read alignment using BWA-MEM (version 0.7.18-r1243-dirty)[39] with default settings. Resulting SAM files were converted to BAM format, sorted, and indexed using SAMtools (version 1.21).

A final round of polishing was performed using Pilon (version 1.24)[40], which incorporated the aligned Illumina reads to correct residual base errors, small insertions and deletions, and local misassemblies. This generated the final polished genome assembly for the ancestral strains.

Assembled genomes were annotated using Bakta (v1.12.0, database version v6.0)[41]. Fosfomycin-selected variants were *de novo* assembled with SPAdes (v3.13.1)[42] using kmer sizes 21, 33, 55, 77, 99 and 127 on --isolate mode.

Assembly statistics of the UTI-34, UTI-59 and the initial assembly of the fosfomycin-selected variants were assessed using CheckM2 (Galaxy v1.1.0)[43]. The average nucleotide alignment of fosfomycin-selected variants with UTI-34 and UTI-59, and therefore confirmation of being derivatives of the correct ancestor, was determined using pyani (v0.3.0-alpha)[44].

### Calling mutations in fosfomycin-selected variants

Mutations in the fosfomycin-selected variants were called by aligning the sequencing reads to the respective annotated ancestor using Snippy (v4.6.0, https://github.com/tseemann/snippy). The fosfomycin-selected variant 59A initially did not have any observed mutations, and therefore, we aligned the assembled genome to the annotated UTI-59 ancestor using Snippy to further identify any mutations present.

### Screening of observed mutations in Escherichia coli genomes isolated from urine

We searched for BakRep[34] database using the following sets of parameters;

*E. coli* genomes isolated from human urine:

“All must match”: Completeness == 100, Species ∼ *Escherichia coli*, Isolation source ∼ Urine, Contamination <= 2.

“At least one must match”: Host ∼ *Homo sapien*, Isolation source ∼ human.

*E. coli* genomes isolated from any urine:

“All must match”: Completeness == 100, Species ∼ *Escherichia coli*, Isolation source ∼ Urine, Contamination <= 2.

The “human urine only” parameters identified 14,165 genomes; however, two records were removed as the genomes were classified as *Escherichia ruysiae* or *Escherichia albertii*. While the “all urine” parameters identified 20,0072 genomes, and six records were removed as the genomes were classified as *Escherichia ruysiae* or *Escherichia albertii*. Therefore, we included a total of 14,163 *E. coli* genomes for the “human urine only” analysis and 20,0066 genomes *E. coli* for the “all urine” analysis.

The ancestor genes with observed mutations in the fosfomycin-selected variants were extracted from the Bakta-annotated UTI-34 and UTI-59 genome (.ffn) and deposited into individual .fasta files. Each ancestral gene CDS were aligned against the 14,163 “human urine only” or 20,066 “all urine” *E. coli* genomes using minimap2 (v2.30-r1287)[37]. Using the PAF coordinates, the best hit regions from each assembly were extracted using Biopython (v1.87)[45] and Python (v3.13.5). Identical CDS sequences were deduplicated using seqkit (v2.13.0)[46]. The deduplicated genes were then cleaned to remove any short (<100 nucleotides), near empty (ungapped length <50% of ancestor CDS) or non-IUPAC sequences using Biopython and Python. Ancestor and clean CDS were aligned using the codon-aware alignment tool, MACSE (v2.07)[47], while preserving the reading frame, and the alignment was cleaned using Biopython and Python to remove near-gap sequences (<50% ungapped). Using the list of observed mutations defined by Snippy, missense mutations were detected by identifying amino acid changes at called positions, frameshift mutations were detected by ! markers in nucleotide alignments at the codon positions, in-frame indels were detected through gaps in amino acid alignments and stop mutations were detected through translated sequences, identifying stop gain and stop loss mutations, all using Biopython and Python. A three-tier evidence system was applied to each mutation type detected within the dataset, with the exact called mutation (exact tier), the same mutation type within ±20 amino acids of the called position (regional tier) and the same mutation type anywhere within the gene (gene level tier). Sequences supporting each confirmed mutation were extracted and the number of unique isolates per mutation was counted. This pipeline was trained on the assembled fosfomycin-selected variants to confirm the accuracy of the called mutations. The scripts for automation of this pipeline were developed in collaboration with large language model AI tools, GPT-5 (OpenAI) and Claude Sonnet 4.6 (Anthropic). AI assistance was used in pipeline design, script development, debugging and code review.

### Fitness

Three biological replicate cultures of UTI-34 and UTI-59 and the fosfomycin-selected variantsw ere diluted to an optical density at 600 nm (OD_600_) of 0.1 in PBS. The diluted culture was then further diluted 1 in 1000 in MH broth, and 100 µl aliquoted to a flat-bottom 96-well plate in duplicate per biological replicate. Negative controls of 100 µl of MH broth were added in duplicate, respectively, and were included per biological replicate. Plates were incubated at 37°C, shaking at 200 rpm for 24 hours in a SPECTROstar NANO microplate reader (BMG Labtech, Germany) with each well read every 10 minutes, 100 flashes per well at OD_600_.

### Statistical analysis

Normality of data was assessed by a Shapiro wilk test. Statistical significance of the disc diffusion data and the growth curve data of the fosfomycin-selected variants was assessed using a one-sample t-test or a one-sample Wilcoxon test to compare the evolved clones against the ancestor. Due to the multiple comparisons, *p*-values were adjusted using the Benjamini-Hochberg method.

## Results

### Minimum inhibitory concentrations in apical urine and selection experiment

We selected UTI-34 and UTI-59 *E. coli* isolates for the evolution experiment due to their different genomic backgrounds, sequence types (ST73 and ST127, respectively[4]), and both isolates were susceptible to antibiotics commonly used to treat UTIs, including fosfomycin[4].

The MIC of fosfomycin against UTI-34 and UTI-59 was assessed in conditioned urine collected from the apical chamber during differentiation of the 3D-UHU from day 10 onwards and MH broth. This ensured that the correct concentration of fosfomycin could be used in the 3D-UHU model, as the environment can impact the activity of an antimicrobial[7]. UTI-34 and UTI-59 were susceptible to fosfomycin according to EUCAST breakpoints in both MH broth and apical urine, and there were no differences between the MIC in the two media (Fig. 1A).

**Figure 1:**
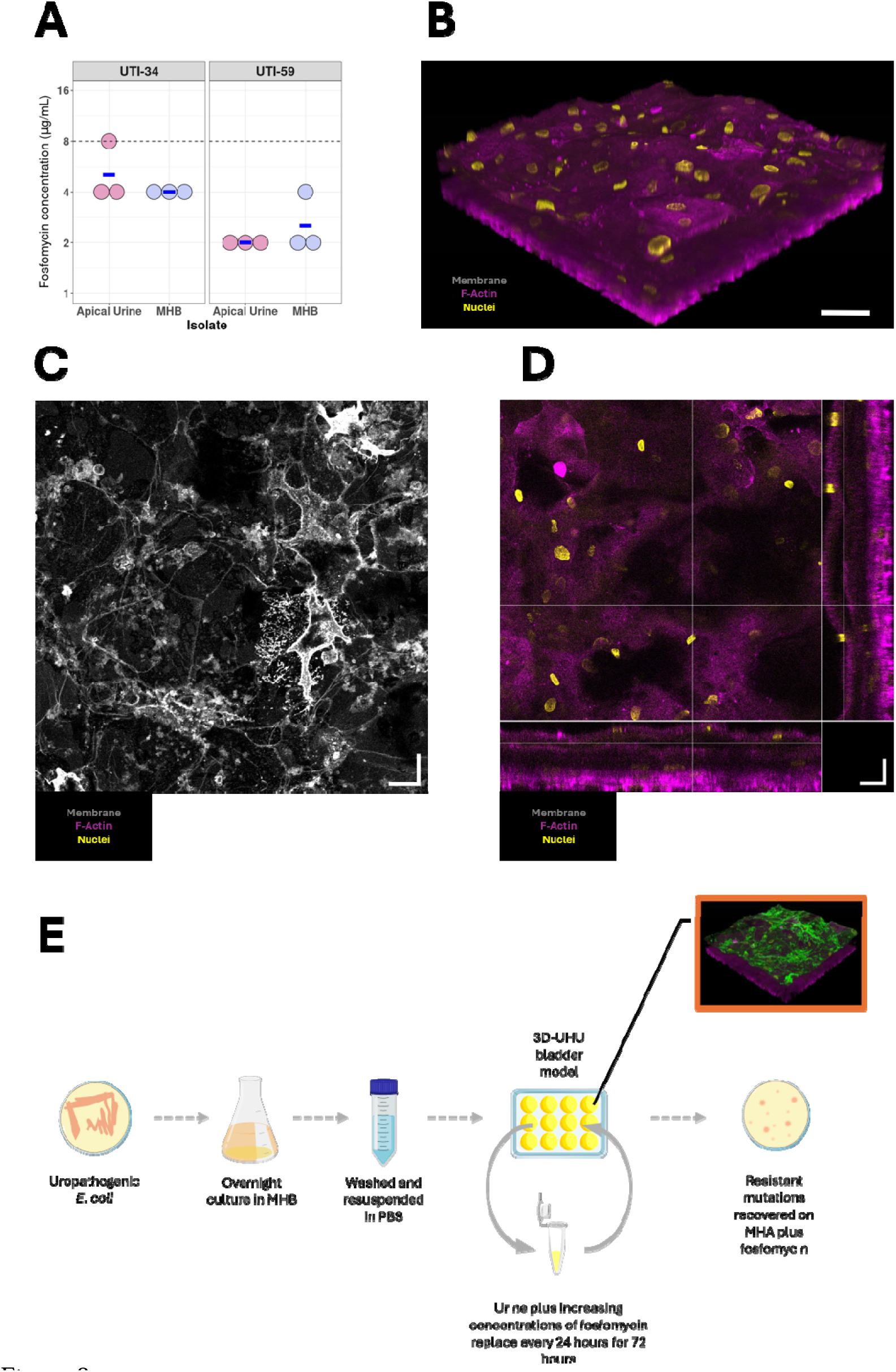
**A)** Minimum inhibitory concentrations of two uropathogenic *Escherichia coli* strains, UTI-34 and UTI-59, in Mueller Hinton broth (MHB) and conditioned apical urine from the 3D urine tolerant human urothelium (3D-UHU) model by broth microdilution method. Confocal imaging of the 3D urine-tolerant human urothelial model in the **B)** top-down (main square) and orthogonal view, **C)** maximum intensity projection and **D)** 3D reconstructed view. Magenta indicates the F-actin or cytoskeleton, yellow indicates cell nuclei and grey indicates the cell membrane of umbrella cells. Scale bar is 20 µm. **E)** Process of experimental evolution of UTI-34 and UTI-59 within the 3D-UHU model. Briefly, UTI-34 and UTI-59 were initially grown for 18 hours in MHB and then washed three times in PBS. The 3D-UHU model was infected with the washed isolates and exposed to 0.25x, 0.5x and 1x MIC of fosfomycin sequentially over 72 hours, where every 24 hours the urine from the 3D-UHU model was removed and fresh urine added with a higher concentration of fosfomycin and re-infected with the bacterial growth from the previous 24 hours. After 72 hours the urine was plated out on to Mueller Hinton agar containing 1x, 2x, 4x and 8x MIC to recover fosfomycin-selected variants of UTI-34 and UTI-59. Created using bioicons (https://bioicons.com/).

The correct 3D structure of the 3D-UHU model was confirmed, with good differentiation of the umbrella cells and good stratification of approximately 30-40 µM [17, 18](Fig. 1B-D and Supplementary Figure S1A and S1B). We performed a modified evolutionary ramp experiment in the 3D-UHU model, with the fosfomycin concentration doubling every 24 hours over the 72 hours of the experiment from 0.25x MIC to 1x MIC. We supplemented the urine in the apical chamber of the 3D-UHU model with 25 µg/ml glucose-6-phosphate to activate UhpT and facilitate the uptake of fosfomycin[48–50]. The evolution experiment was performed in 5% CO_2,_ and the variants were recovered on MHA containing 4x and 8x MIC of fosfomycin (Fig. 1D). We generated five independently evolved variants from each ancestor.

### Mutational landscape of fosfomycin-selected variants

The variant 34D was removed from further analysis as it was found to be a contaminant following sequencing (Supplementary Table S1 and Supplementary Figure S2). We identified mutations in the fosfomycin-selected variants bioinformatically using Snippy and alignment with the respective ancestor genome. Diverse mutations were observed in UTI-34 fosfomycin-selected variants. Both 34A and 34B contained deletions of different sizes and positions in *glpT*, resulting in a conservative in-frame Trp256 to Leu268 deletion and a disruptive in-frame Trp182 to His187 deletion with a cysteine insertion, respectively (Fig. 2A and Supplementary Table S2). Variant 34C harboured a single nucleotide polymorphism (SNP) in *glpT* leading to an Ala164Val missense (Fig. 2A and Supplementary Table S2). In addition to the deletion in *glpT*, a SNP in *uhpC* was also identified in 34B, resulting in a loss of the stop codon at Ter440Cysext* (Fig. 2A and Supplementary Table S2). Three separate mutations were found in 34E; a SNP in *uhpT* causing a nonsense mutation at Gln7*, a SNP in *cyaA* resulting in a missense at Tyr394Asp, and a single base pair insertion in *ptsI* causing a frameshift at Ala294fs (Fig. 2A and Supplementary Table S2). In contrast, UTI-59 fosfomycin-selected variants were less diverse. A single base pair deletion in both *uhpT* and *glpT* was found in all UTI-59 fosfomycin-selected variants, except 59D, which led to Ala423fs and Ala294fs frameshifts, respectively (Fig. 2B and Supplementary Table S2). Instead, 59D contained a 9 bp deletion in *uhpA,* resulting in a conservative in-frame deletion at Ala41 to Leu43 (Fig. 2B and Supplementary Table S2). Off-target mutations were also identified in 34A, with an intragenic SNP, and 59C and 59E containing SNPs or complex mutations in *tnpA*, an IS66-like element accessory protein.

**Figure 2:**
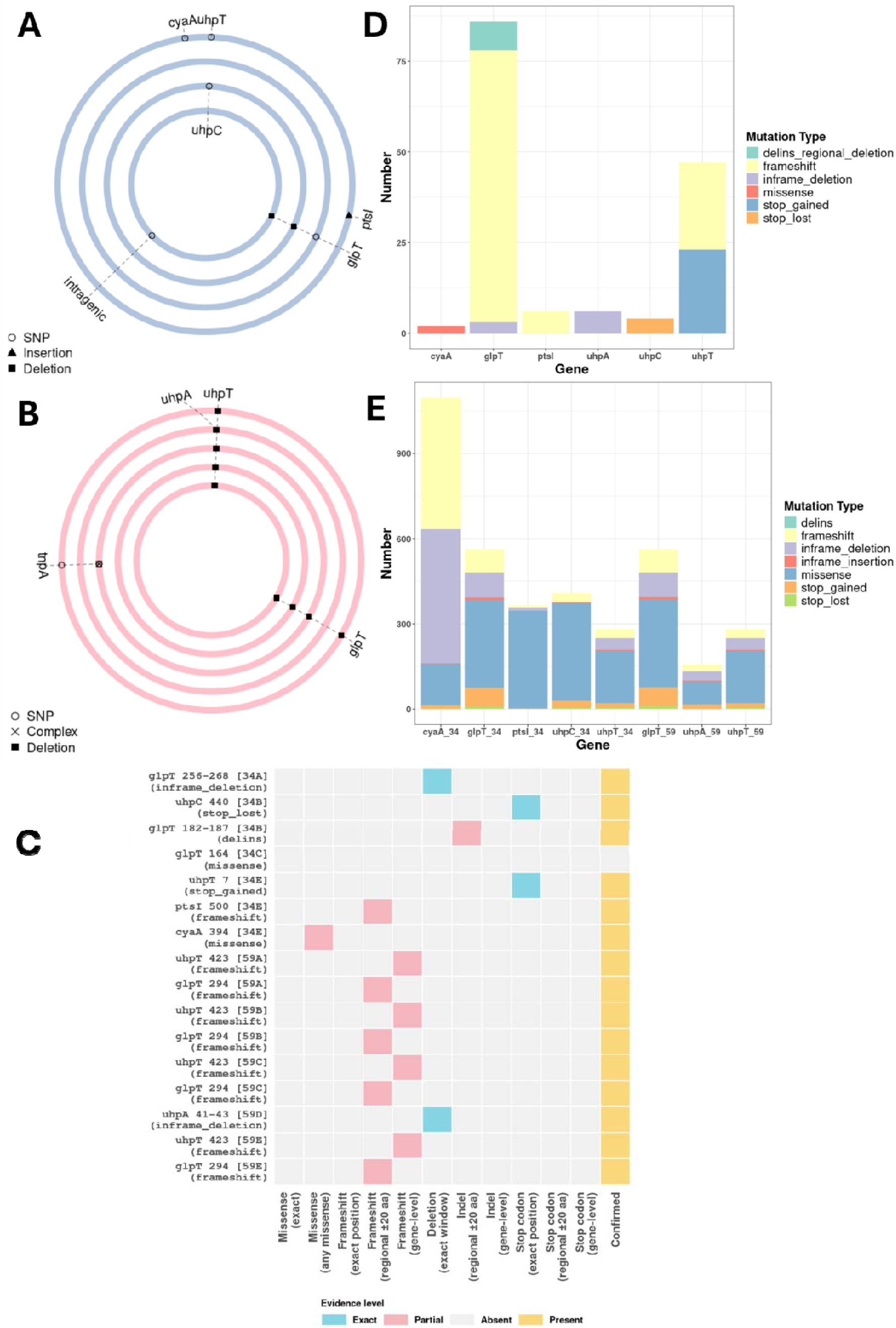
Relative positions of mutations in fosfomycin-selected variants of **A)** UTI-34 and **B)** UTI-59 as identified by Snippy. **C)** Presence or absence of the amino acid changes found the in the fosfomycin-selected variants of UTI-34 and UTI-59 in the “human urine only” 14,163 *E. coli* genomes isolated from human urine. Exact tier evidence identifies the exact expected mutation found in the fosfomycin-selected variants whereas any missense is a missense at the exact position but a different amino acid change than expected. Regional tier evidence calls the same mutation type 20 amino acids up and downstream of the expected position and gene tier calls the same mutation type anywhere in the gene. **D)** The number of unique genomes with the called tiered mutation within the “human urine only” genome dataset per gene. **E)** The spectrum and number of the different mutation types found in each gene relative the corresponding gene found in the ancestralUTI-34 and UTI-59.

### Screening against the BakRep libraries

To confirm whether the mutations we have identified in *glpT*, *uhpT*, uhpA, uhpC, *ptsI* and *cyaA* are found in *E. coli* isolates isolated from urine, we developed a pipeline to screen a dataset for these mutations. A dataset of *E. coli* genomes from all urine and human urine from BakRep[34], a database of genomes which have been consistently assembled and are deposited with searchable metadata, was assembled. While the *E. coli* genomes found in BakRep represent a robust dataset of circulating *E. coli*, they were not specifically fosfomycin-resistant isolates and, therefore, we expected limited representation of diverse fosfomycin-resistant mutations in the dataset. For this reason, we applied a tiered approach to identifying the mutations found in our fosfomycin-selected variants within this dataset. Specifically, we screened for the exact mutation (exact mutation type at the exact position), regional mutation (exact mutation type within ±20 amino acids of the expected position) and gene mutation (exact mutation type anywhere within the gene). We first validated the pipeline by screening the mutations called by Snippy against the fosfomycin-selected variant genomes. We confirmed the exact frameshift mutation in *glpT* and *uhpT*, in-frame deletion in *glpT* and *uhpA*, loss of the stop codon in *uhpC*, nonsense mutation in *uhpT* and missense in *cyaA* and *glpT* (Supplementary Figure S4). The frameshifts in *ptsI* and *glpT* were found at the regional tier. However, manual inspection of the extracted sequences containing the evidence of these mutations from the genome identified the exact presence of these mutations (Supplementary Figure S4). Therefore, all mutations called by Snippy were confirmed to be present in the genomes using the pipeline.

We next screened the mutations identified in our fosfomycin-selected variants against the “all urine” dataset of 20,066 *E. coli* genomes. This dataset predominantly included genomes of *E. coli* isolated from human, canine and feline urine. We confirmed the presence of the exact in-frame deletion in *glpT* and *uhpA*, the exact loss of the stop codon in *uhpC* and the nonsense mutation in *uhpT* (Supplementary Figure S5A). While we did not observe the exact missense amino acid change in *cyaA*, we did find an alternative missense at the same position (Supplementary Figure S5A). All frameshift mutations were identified at the regional tier, within 20 amino acids upstream and downstream of the expected mutation site, in the “all urine dataset” (Supplementary Figure S5A). The exact deletion in *glpT* was present but without the insertion of cysteine, and we did not observe any missense mutations at position 164 in *glpT*. The incidence of unique confirmed evidence of these mutations within the “all urine” dataset varied per gene. Any missense mutations at position 394 in *cyaA* (2), in-frame deletions and region deletion-insertions in *glpT* (3 and 10, respectively), regional frameshifts in *ptsI* (9) and loss of the stop codon in *uhpC* (6) were few. In contrast, regional frameshifts in *glpT* (94) and *uhpT* (28), in-frame deletions in *uhpA* (28) and *uhpT* (25) and nonsense mutations were more common.

As we were using a model which mimicked the human urothelium, we further refined the *E. coli* genome dataset to screen for our observed mutations in genomes isolated only from human urine, “human urine only”. Again, the exact in-frame deletion in *glpT* and *uhpA*, loss of the stop codon in *uhpC* and the nonsense mutation in *uhpT* were all confirmed, as were the regional frameshifts in *ptsI* and *glpT*, alternative missense at position 394 in *cyaA* and regional deletion in *glpT* without the cysteine insertion (Figure 2C). In fact, the only difference between the “all urine” and “human urine only” dataset was that *uhpT* frameshifts were only found at a gene level rather than at the regional level (Figure 2C). Again, the incidence of unique confirmed evidence of these mutations within the “human urine only” dataset varied per gene. There was low incidence of any missense mutations at position 394 in *cyaA* (2), in-frame deletions in *glpT* (3) and *uhpA* (6), regional deletion-insertions in *glpT* (8), frameshifts in *ptsI* (6) and loss of the stop codon in *uhpC* (4) (Fig. 2D). In contrast, there were relatively high evidence of the regional frameshifts in *glpT* (75) and gene-level frameshift (24) and early termination (23) in *uhpT* (Fig. 2D). In general, frameshifts, in-frame deletions, nonsense and missense mutations are represented in all six genes, although to varying degrees (Figure 2E). In-frame deletions and stop-lost mutations were present in all but *ptsI* (Figure 2E).

### Collateral susceptibility and fitness

All fosfomycin-selected variants were below the EUCAST clinical breakpoint via disc diffusion, confirming resistance to fosfomycin in all fosfomycin-selected variants (UTI-34 *p*-value = 0.00029 UTI-59 *p*-value = 0.0098, Fig. 3A and 3B). There were no observed statistically significant increases or decreases in susceptibility of the fosfomycin-selected variants to cephalexin (UTI-34 *p*-value = 0.31; UTI-59 *p*-value = 0.31) and nitrofurantoin (UTI-34 *p*-value = 0.31; UTI-59 *p*-value = 0.72) relative to UTI-34 and UTI-59. For trimethoprim, UTI-34 had a statistically significant increase in susceptibility (*p*-value = 0.012), but this was not observed in UTI-59 (*p*-value=0.41) (Fig. 3A and 3B). This suggests that these mutations do not cause collateral susceptibility or collateral resistance to these antibiotics when assessed by disc diffusion on MH agar.

**Figure 3:**
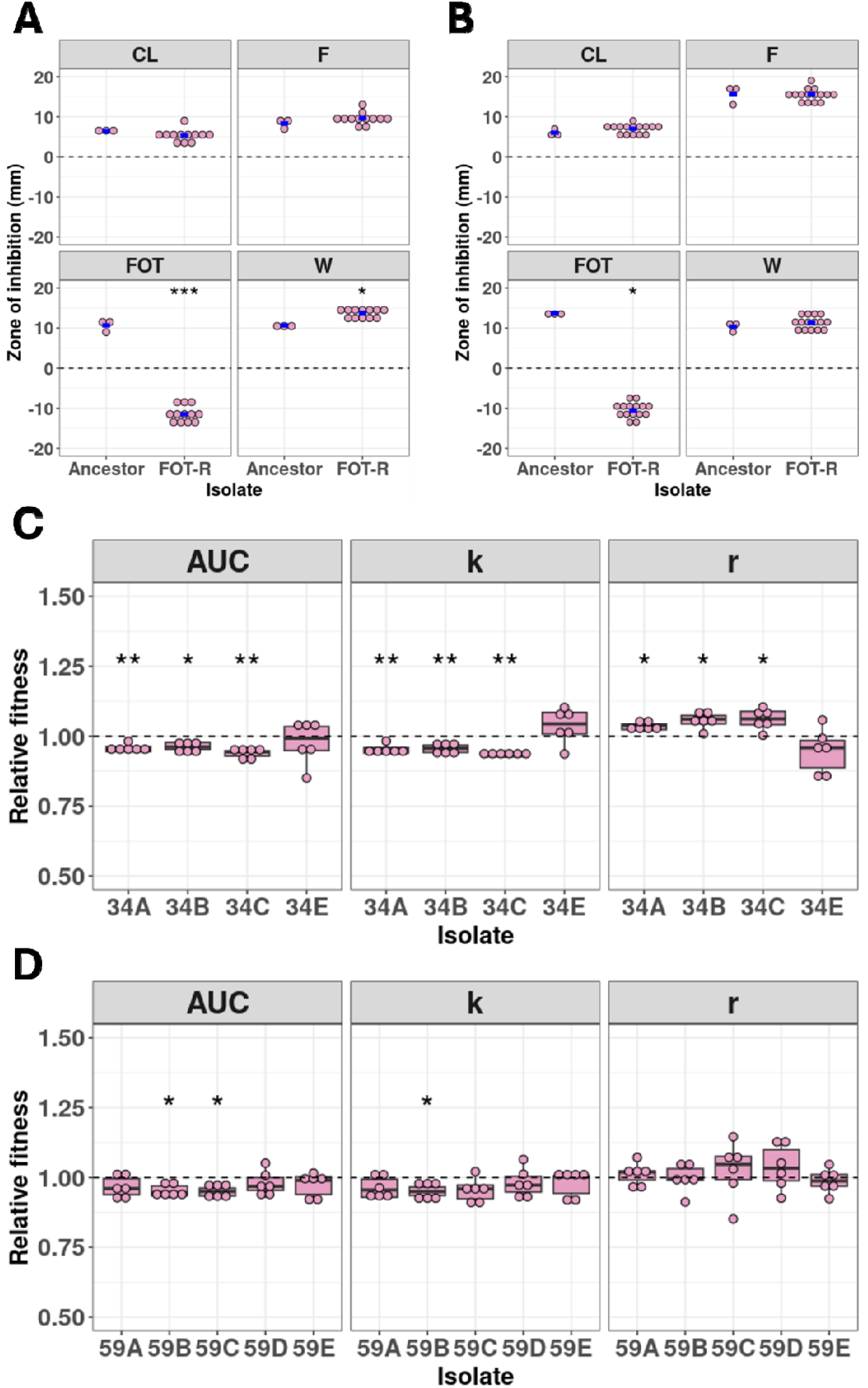
Collateral susceptibility of **A)** UTI-34 and **B)** UTI-59 (ancestors) and the fosfomycin-selected variants (FOT-R) against cephalexin (CL), nitrofurantoin (F), fosfomycin (FOT) and trimethoprim (F), assessed by disc diffusion. Fitness of fosfomycin-selected variants of C) UTI-34 and D) UTI-59 in Mueller Hinton broth relative to the respective ancestors. Fitness was determined by the parameters, area under the curve (AUC), carrying capacity (*k*) and maximum growth rate (*r*).

Fitness of the fosfomycin-selected variants were assessed by growth curves in MH broth in the absence of fosfomycin. In UTI-34 fosfomycin-selected variants, we observed a significant decrease in area under the curve (AUC, *p*-value = 0.0048-0.014) and carrying capacity (*k*, *p*-value = 0.0011-0.0086) and a significant increase in maximum growth rate (*r*, *p*-value = 0.013-0.025) relative to UTI-34 in MH broth in all variants except 34E (Fig. 3C). In 34E, there was no significant difference in AUC (*p*-value =0.6), k (*p*-value = 0.31) and *r* (*p*-value = 0.27) relative to UTI-34 in MH broth (Fig. 3C). In contrast, for fosfomycin-selected variants of UTI-59, most variants had no significant difference in AUC (*p*-value = 0.15-0.41), *k* (*p*-value = 0.11-1.0) and *r* (*p*-value = 0.6-0.93) relative to UTI-59 in MH broth (Fig. 3D). Variant 59B had a significant decrease in AUC (*p*-value = 0.013) and in *k* (*p*-value = 0.018) compared to UTI-59 in MH broth (Fig 3D), while variant 59C also had a significant decrease in AUC (*p*-value 0.013) compared to UTI-59 (Fig 3D).

## Discussion

The clinical relevance of *in vitro* studies on AMR has been limited, at least partially, due to the lack of correlation between laboratory growth media and the *in vivo* environment. Fundamental understanding of AMR from *in vitro* axenic studies is essential, but to fully explore and combat AMR we must study AMR in the context in which it exists, for example a more relevant infection environment. In this proof-of-concept study, we have demonstrated the selection of clinically relevant mutations in two uropathogenic *E. coli* clinical isolates which confer resistance to fosfomycin in the 3D-UHU model, which closely replicates the human bladder urothelium. Diverse mutations were selected for and all fosfomycin-selected variants contained mutations in genes previously linked to clinical fosfomycin resistance[29–33]. This novel approach has the potential to enable prospective studies of AMR acquisition under treatment, and the ability to manipulate the urine used in the model, for example to correspond to specific patient cohorts, different antimicrobial combinations or include mono- and polymicrobial infections, is an attractive property of 3D-UHU model.

To confirm clinical relevance of the observed fosfomycin-selected mutation, we screened these mutations against a dataset of clinical isolates of *E. coli* isolated from urine. Although matched phenotype and genotype data are available, currently there are limitations in AMR prediction tools to predict loss-of-function mutations[51, 52], many of which are represented in the fosfomycin-selected variants. Matched genotype-phenotype datasets could therefore bias conclusions by limiting the number of genomes to only those which contain the relatively few fosfomycin-resistant mutations which can be called by the AMR prediction tools. An alternative approach is using BakRep[34], a database which provides high-quality, consistently assembled genomes and searchable metadata. Using the BakRep database, we built an “all urine” and “human urine only” *E. coli* genome dataset to screen the mutations observed in the fosfomycin-selected variants. Isolates selected for whole genome sequencing are more likely to be antimicrobial resistant, thereby biasing the genomes in BakRep. While this may increase the proportion of *E. coli* in the BakRep database which are fosfomycin resistant above what is seen nationally and globally, we still expected there to be a limited number with clinically relevant mutations in the target genes. One limitation of this database is the inability to confirm that these identified mutations cause fosfomycin resistance. However, as we are screening the known mutations from the fosfomycin-selected variants, which have been confirmed to cause fosfomycin resistance, this could be inferred. Therefore, we determined that the limitation of being unable to match genotype to phenotype was offset by having an unbiased but specific dataset.

GlpT and UhpT are both antiporters which facilitate the exchange of a phosphate ion to allow the uptake of glycerol-3-phosphate and glucose-6-phosphate, respectively[48–50]. As fosfomycin is a phosphate-based antibiotic, GlpT and UhpT both facilitate the transport of fosfomycin into the cell[48–50]. The expression of UhpT is induced by glucose-6-phospahte by interacting with the UhpT regulator UhpABC. UhpC and UhpB are the sensor-histidine kinase two-component system which activates the transcriptional regulatory protein, UhpA[50]. The expression of GlpT and UhpT is reliant on cyclic AMP (cAMP), which upregulates their expression[48–50]. The formation of cAMP from ATP is catalysed by CyaA, activated by glucose-6-phosphate and the carbohydrate phosphotransferase system (PTS), which is formed of PtsI, PtsH and Enzyme II[50]. Once in the cell, fosfomycin inhibits the first committed step of peptidoglycan synthesis by covalently binding to MurA, preventing the formation of UDP-*N*-acetylglucosamine-enolpyruvyltransferase[48–50].

All fosfomycin-selected variants contained a mutation in *glpT*, *uhpT*, *uhpA* and/or *uhpC*. Mataseje *et al*. found that 57 of 5012 isolates of *E. coli* from the CANWARD surveillance programme from 2007 to 2022 were fosfomycin-resistant or fosfomycin-intermediate[32]. Within these 57 *E. coli* isolates, disruption to GlpT and UhpT transporters and the UhpABC regulatory systems were common, with frameshift and deletions found in *uhpT*, *uhpA* and *uhpC* and missense and nonsense identified in *glpT*, *uhpT*, *uhpA* and *uhpC*[32]. Insertion and/or deletions linked to fosfomycin resistance have also been previously found in *uhpT*, *glpT* and *uhpA[53, 54]*. While missense mutations have been observed in *ptsI* and *cyaA,* and nonsense mutations observed in cyaA, it was unclear if they contributed to fosfomycin resistance[32, 50]. Importantly, mutations in *ptsI* and *cyaA* in the fosfomycin-selected variants only occurred in 34E and alongside a nonsense mutation in *uhpT*.

While this study has produced promising data, there are limitations which need to be explored further in larger studies. While the two clinical isolates of *E. coli* had different sequence types and distinct AMR profiles, this represents a limited panel. Genetic background can affect the selection of mutations in bacteria[55, 56] and therefore the panel of isolates used should be expanded to include diverse sequence types of *E. coli* and other uropathogens. Indeed, other antimicrobials, where resistance is driven at least partially by mutation, used for the treatment of UTIs should also be explored within the 3D-UHU model beyond fosfomycin to determine if clinically relevant mutations are also selected for. Additionally, AMR can be horizontally transferred by mobile genetic elements in artificial urine[57] and therefore it would be important to assess if mobile genetic elements can be transferred within the 3D-UHU model.

Like many other infection sites, the urogenital tract is known to contain a microbiome[58], although it is often simpler and of a lower biomass than other sites, such as the gut[58]. While the healthy female urogenital microbiome is dominated by *Lactobacillus* spp.[58], a disrupted microbiome has reduced *Lactobacillus*, is more diverse and has been thought to potentiate UTIs and recurrent UTIs[58–61]. It has been demonstrated that the presence of other bacteria can impact the acquisition of AMR[57]. In this study we have only used a monoculture to assess AMR acquisition, but the role of the healthy and disrupted urogenital microbiome on AMR acquisition would be important to explore. Despite these limitations, this study has shown that the 3D-UHU model could be used to further explore AMR acquisition in more physiologically relevant environments and has the potential to produce clinically relevant data. These data could help better understand AMR in an infection context, optimise antimicrobial regimens and develop novel non-antimicrobial treatments, all to help limit the acquisition of AMR during UTI therapy.

## Conclusions

In this study we have used the 3D-UHU model which closely mimics the human bladder urothelium to select for fosfomycin-resistant variants of two uropathogenic *E. coli* strains. By screening these observed mutations against a curated but unbiased dataset of *E. coli* genomes isolated from urine, we confirmed that the mutations observed in the fosfomycin-selected variants were representative of those found in clinical isolates of *E. coli*. The use of 3D culture models which more closely recapitulate the human infection environment could improve the understanding of how AMR evolves, is acquired and how it is expressed *in vivo*. Additionally, increasing the clinical relevance of *in vitro* AMR studies could inform new strategies to combat AMR.

## Author Contributions

B.J. – Investigation, Methodology, Resources, Validation and Writing – reviewing & editing; M.W. – Investigation, Methodology, Resources and Writing – reviewing & editing; M.F. – Methodology, Resources, Validation and Writing – reviewing & editing; B.O.M. – Methodology, Resources, Validation and Writing – reviewing & editing; D.J.W. – Investigation, Methodology, Resources and Writing – reviewing & editing; C.C. – Formal Analysis and Writing – reviewing & editing; J.R. – Methodology, Resources and Writing – reviewing & editing; A.T.M.H. – Conceptualization, Data Curation, Formal Analysis, Funding Acquisition, Investigation, Methodology, Project Administration, Software, Supervision, Validation, Visualization and Writing – original draft.

## Acknowledgements

We would like to thank Dr Conor Meehan, Nottingham Trent University, for their support with statistical analysis of the MIC, disc diffusion and fitness data and Dr Graham Hickman, Nottingham Trent University, for his support with confocal imaging.

## Data availability

Illumina and Oxford Nanopore Technologies sequencing reads generated for UTI-34 and UTI-59 can be found in the sequencing read archive (SRA) under BioProject number PRJNA1037559. All sequencing reads generated during this project are available in the sequencing read archive (SRA) under the BioProject number PRJNA1491657. All commands and code for bioinformatics analysis, generation of figures and statistical analysis, bakrep-export files, key outputs, fasta files and datasets for figures are available at https://github.com/Alhubb/fos-selection-in-3D-UHU.

## References

1. Cazares D, Rayner E, Cazares A, Figueroa W, Goulding D, et al. Eco-evolutionary responses to plasmid-dependent phage constrain the spread of multidrug-resistance plasmids. ISME J;20. Epub ahead of print 14 January 2026. DOI: 10.1093/ismejo/wrag113.

2. Jago MJ, Soley JK, Denisov S, Walsh CJ, Gifford DR, et al. High-throughput method characterizes hundreds of previously unknown antibiotic resistance mutations. Nat Commun 2025;16:780.

3. Hubbard ATM, Mason J, Roberts P, Parry CM, Corless C, et al. Piperacillin/tazobactam resistance in a clinical isolate of Escherichia coli due to IS26-mediated amplification of blaTEM-1B. Nat Commun 2020;11:4915.

4. James B, Reesaul H, Kashif S, Behruznia M, Meehan CJ, et al. The effect of antibiotic selection on collateral effects and evolvability of uropathogenic Escherichia coli. npj Antimicrobials and Resistance 2024;2:19.

5. Eladawy M, McCartney AL, Garner AC, Hoyles L. Growth of Gram-negative uropathogens in human urine under hypoxic conditions produces clinically relevant metabolomic profiles. BioRxiv. Epub ahead of print 7 May 2025. DOI: 10.1101/2025.05.07.652693.

6. Hinz A, Amado A, Kassen R, Bank C, Wong A. Unpredictability of the Fitness Effects of Antimicrobial Resistance Mutations Across Environments in *Escherichia coli*. Mol Biol Evol;41. Epub ahead of print 3 May 2024. DOI: 10.1093/molbev/msae086.

7. Hubbard ATM, Jafari N V., Feasey N, Rohn JL, Roberts AP. Effect of Environment on the Evolutionary Trajectories and Growth Characteristics of Antibiotic-Resistant Escherichia coli Mutants. Front Microbiol;10. Epub ahead of print 28 August 2019. DOI: 10.3389/fmicb.2019.02001.

8. Garcia-Maset R, Chu V, Yuen N, Blumgart D, Yoon J, et al. Effect of host microenvironment and bacterial lifestyles on antimicrobial sensitivity and implications for susceptibility testing. npj Antimicrobials and Resistance 2025;3:42.

9. Shepherd MJ, Harrington NE, Kotarra A, Igler C, Cagney K, et al. Multiple ecological and evolutionary mechanisms drive treatment-induced antibiotic resistance. BioRxiv. Epub ahead of print 19 February 2026. DOI: 10.64898/2026.02.18.705331.

10. Diaz Caballero J, Wheatley RM, Kapel N, López-Causapé C, Van der Schalk T, et al. Mixed strain pathogen populations accelerate the evolution of antibiotic resistance in patients. Nat Commun 2023;14:4083.

11. DelaFuente J, Toribio-Celestino L, Santos-Lopez A, León-Sampedro R, Alonso-del Valle A, et al. Within-patient evolution of plasmid-mediated antimicrobial resistance. Nat Ecol Evol 2022;6:1980–1991.

12. Doroshenko N, Rimmer S, Hoskins R, Garg P, Swift T, et al. Antibiotic functionalised polymers reduce bacterial biofilm and bioburden in a simulated infection of the cornea. Biomater Sci 2018;6:2101–2109.

13. Okurowska K, MacNeil S, Roy S, Garg P, Monk PN, et al. Exploring interspecies differences in ex vivo models of Pseudomonas aeruginosa keratitis: a comparative study of human, pig and sheep corneas. J Med Microbiol;73. Epub ahead of print 13 December 2024. DOI: 10.1099/jmm.0.001901.

14. Jordana-Lluch E, Garcia V, Kingdon ADH, Singh N, Alexander C, et al. A Simple Polymicrobial Biofilm Keratinocyte Colonization Model for Exploring Interactions Between Commensals, Pathogens and Antimicrobials. Front Microbiol;11. Epub ahead of print 26 February 2020. DOI: 10.3389/fmicb.2020.00291.

15. Harrington NE, Sweeney E, Harrison F. Building a better biofilm - Formation of in vivo-like biofilm structures by Pseudomonas aeruginosa in a porcine model of cystic fibrosis lung infection. Biofilm 2020;2:100024.

16. Harrison F, Muruli A, Higgins S, Diggle SP. Development of an *Ex Vivo* Porcine Lung Model for Studying Growth, Virulence, and Signaling of Pseudomonas aeruginosa. Infect Immun 2014;82:3312–3323.

17. Jafari N V., Rohn JL. An immunoresponsive three-dimensional urine-tolerant human urothelial model to study urinary tract infection. Front Cell Infect Microbiol;13. Epub ahead of print 27 March 2023. DOI: 10.3389/fcimb.2023.1128132.

18. Flores C, Ling J, Loh A, Maset RG, Aw A, et al. A human urothelial microtissue model reveals shared colonization and survival strategies between uropathogens and commensals. Sci Adv;9. Epub ahead of print 10 November 2023. DOI: 10.1126/sciadv.adi9834.

19. National Institute for Health and Care Excellence. Urinary tract infection (lower): antimicrobial prescribing. https://www.nice.org.uk/guidance/ng109 (31 October 2018, accessed 22 June 2026).

20. Kettlewell R, Jones C, Felton TW, Lagator M, Gifford DR. Insights into durability against resistance from the antibiotic nitrofurantoin. npj Antimicrobials and Resistance 2024;2:41.

21. UK Health Security Agency. Antimicrobial Resistance in Escherichia coli bacteriuria. https://ukhsa-dashboard.data.gov.uk/antimicrobial-resistance/antimicrobial-resistance-in-e-coli-bacteriuria (28 May 2026, accessed 22 June 2026).

22. UK Health Security Agency. English surveillance programme for antimicrobial utilisation and resistance (ESPAUR). London; 2025.

23. Osei Sekyere J. Genomic Insights Into Nitrofurantoin Resistance Mechanisms and Epidemiology in Clinical Enterobacteriaceae. Future Sci OA;4. Epub ahead of print 27 June 2018. DOI: 10.4155/fsoa-2017-0156.

24. Wan Y, Mills E, Leung RCY, Vieira A, Zhi X, et al. Alterations in chromosomal genes nfsA, nfsB, and ribE are associated with nitrofurantoin resistance in Escherichia coli from the United Kingdom. Microb Genom;7. Epub ahead of print 24 December 2021. DOI: 10.1099/mgen.0.000702.

25. Kettlewell R, Jones C, Felton TW, Lagator M, Gifford DR. Insights into durability against resistance from the antibiotic nitrofurantoin. npj Antimicrobials and Resistance 2024;2:41.

26. Somorin YM, Weir N-JM, Pattison SH, Crockard MA, Hughes CM, et al. Antimicrobial resistance in urinary pathogens and culture-independent detection of trimethoprim resistance in urine from patients with urinary tract infection. BMC Microbiol 2022;22:144.

27. Grilo T, Freire S, Miguel B, Martins LN, Menezes MF, et al. Occurrence of plasmid-mediated fosfomycin resistance (fos genes) among Escherichia coli isolates, Portugal. J Glob Antimicrob Resist 2023;35:342–346.

28. Abdelraheem WM, Mahdi WKM, Abuelela IS, Hassuna NA. High incidence of fosfomycin-resistant uropathogenic E. coli among children. BMC Infect Dis 2023;23:475.

29. Li Y, Zheng B, Li Y, Zhu S, Xue F, et al. Antimicrobial Susceptibility and Molecular Mechanisms of Fosfomycin Resistance in Clinical Escherichia coli Isolates in Mainland China. PLoS One 2015;10:e0135269.

30. Cottell JL, Webber MA. Experiences in fosfomycin susceptibility testing and resistance mechanism determination in Escherichia coli from urinary tract infections in the UK. J Med Microbiol 2019;68:161–168.

31. Campos AC da C, Andrade NL, Couto N, Mutters NT, de Vos M, et al. Characterization of fosfomycin heteroresistance among multidrug-resistant Escherichia coli isolates from hospitalized patients in Rio de Janeiro, Brazil. J Glob Antimicrob Resist 2020;22:584–593.

32. Mataseje LF, Lysak C, Lerminiaux N, Baxter M, Karlowsky JA, et al. Mechanisms of fosfomycin resistance observed in clinical isolates of *Escherichia coli* from Canada: CANWARD 2007–2022. JAC Antimicrob Resist;8. Epub ahead of print 7 January 2026. DOI: 10.1093/jacamr/dlag020.

33. Nilsson AI, Berg OG, Aspevall O, Kahlmeter G, Andersson DI. Biological Costs and Mechanisms of Fosfomycin Resistance in *Escherichia coli*. Antimicrob Agents Chemother 2003;47:2850–2858.

34. Fenske L, Jelonek L, Goesmann A, Schwengers O. BakRep – a searchable large-scale web repository for bacterial genomes, characterizations and metadata. Microb Genom;10. Epub ahead of print 30 October 2024. DOI: 10.1099/mgen.0.001305.

35. Bolger AM, Lohse M, Usadel B. Trimmomatic: a flexible trimmer for Illumina sequence data. Bioinformatics 2014;30:2114–2120.

36. Wick RR, Howden BP, Stinear TP. Autocycler: long-read consensus assembly for bacterial genomes. Bioinformatics;41. Epub ahead of print 1 September 2025. DOI: 10.1093/bioinformatics/btaf474.

37. Li H. Minimap2: pairwise alignment for nucleotide sequences. Bioinformatics 2018;34:3094–3100.

38. Vaser R, Sović I, Nagarajan N, Šikić M. Fast and accurate de novo genome assembly from long uncorrected reads. Genome Res 2017;27:737–746.

39. Li H, Durbin R. Fast and accurate short read alignment with Burrows–Wheeler transform. Bioinformatics 2009;25:1754–1760.

40. Walker BJ, Abeel T, Shea T, Priest M, Abouelliel A, et al. Pilon: An Integrated Tool for Comprehensive Microbial Variant Detection and Genome Assembly Improvement. PLoS One 2014;9:e112963.

41. Schwengers O, Jelonek L, Dieckmann MA, Beyvers S, Blom J, et al. Bakta: rapid and standardized annotation of bacterial genomes via alignment-free sequence identification. Microb Genom;7. Epub ahead of print 30 November 2021. DOI: 10.1099/mgen.0.000685.

42. Bankevich A, Nurk S, Antipov D, Gurevich AA, Dvorkin M, et al. SPAdes: A New Genome Assembly Algorithm and Its Applications to Single-Cell Sequencing. Journal of Computational Biology 2012;19:455–477.

43. Chklovski A, Parks DH, Woodcroft BJ, Tyson GW. CheckM2: a rapid, scalable and accurate tool for assessing microbial genome quality using machine learning. Nat Methods 2023;20:1203–1212.

44. Pritchard L, Glover RH, Humphris S, Elphinstone JG, Toth IK. Genomics and taxonomy in diagnostics for food security: soft-rotting enterobacterial plant pathogens. Analytical Methods 2016;8:12–24.

45. Cock PJA, Antao T, Chang JT, Chapman BA, Cox CJ, et al. Biopython: freely available Python tools for computational molecular biology and bioinformatics. Bioinformatics 2009;25:1422–1423.

46. Shen W, Le S, Li Y, Hu F. SeqKit: A Cross-Platform and Ultrafast Toolkit for FASTA/Q File Manipulation. PLoS One 2016;11:e0163962.

47. Ranwez V, Douzery EJP, Cambon C, Chantret N, Delsuc F. MACSE v2: Toolkit for the Alignment of Coding Sequences Accounting for Frameshifts and Stop Codons. Mol Biol Evol 2018;35:2582–2584.

48. Falagas ME, Vouloumanou EK, Samonis G, Vardakas KZ. Fosfomycin. Clin Microbiol Rev 2016;29:321–347.

49. Silver LL. Fosfomycin: Mechanism and Resistance. Cold Spring Harb Perspect Med 2017;7:a025262.

50. Mattioni Marchetti V, Hrabak J, Bitar I. Fosfomycin resistance mechanisms in Enterobacterales: an increasing threat. Front Cell Infect Microbiol;13. Epub ahead of print 4 July 2023. DOI: 10.3389/fcimb.2023.1178547.

51. Madden DE, Webb JR, Steinig EJ, Currie BJ, Price EP, et al. Taking the next-gen step: Comprehensive antimicrobial resistance detection from Burkholderia pseudomallei. EBioMedicine 2021;63:103152.

52. Madden DE, Baird T, Bell SC, McCarthy KL, Price EP, et al. Keeping up with the pathogens: improved antimicrobial resistance detection and prediction from Pseudomonas aeruginosa genomes. Genome Med 2024;16:78.

53. Takahata S, Ida T, Hiraishi T, Sakakibara S, Maebashi K, et al. Molecular mechanisms of fosfomycin resistance in clinical isolates of Escherichia coli. Int J Antimicrob Agents 2010;35:333–337.

54. Ohkoshi Y, Sato T, Suzuki Y, Yamamoto S, Shiraishi T, et al. Mechanism of Reduced Susceptibility to Fosfomycin in *Escherichia coli* Clinical Isolates. Biomed Res Int 2017;2017:1–8.

55. Santos-Lopez A, Marshall CW, Haas AL, Turner C, Rasero J, et al. The roles of history, chance, and natural selection in the evolution of antibiotic resistance. Elife;10. Epub ahead of print 25 August 2021. DOI: 10.7554/eLife.70676.

56. Hoeksema M, Jonker MJ, Brul S, ter Kuile BH. Effects of a previously selected antibiotic resistance on mutations acquired during development of a second resistance in Escherichia coli. BMC Genomics 2019;20:284.

57. Bustamante M, Koopman F, Martens J, Brons JK, DelaFuente J, et al. Community context influences the conjugation efficiency of *Escherichia coli*. FEMS Microbes;5. Epub ahead of print 10 January 2024. DOI: 10.1093/femsmc/xtae023.

58. Neugent ML, Kumar A, Hulyalkar N V., Lutz KC, Nguyen VH, et al. Recurrent urinary tract infection and estrogen shape the taxonomic ecology and function of the postmenopausal urogenital microbiome. Cell Rep Med 2022;3:100753.

59. Bossa L, Kline K, McDougald D, Lee BB, Rice SA. Urinary catheter-associated microbiota change in accordance with treatment and infection status. PLoS One 2017;12:e0177633.

60. Meštrović T, Matijašić M, Perić M, Čipčić Paljetak H, Barešić A, et al. The Role of Gut, Vaginal, and Urinary Microbiome in Urinary Tract Infections: From Bench to Bedside. Diagnostics 2020;11:7.

61. Thomas-White KJ, Gao X, Lin H, Fok CS, Ghanayem K, et al. Urinary microbes and postoperative urinary tract infection risk in urogynecologic surgical patients. Int Urogynecol J 2018;29:1797–1805.

