## Supplementary Data for "*In vitro* evolution of uropathogenic *Escherichia coli* to fosfomycin resistance in a 3D cultured human bladder microtissue model"

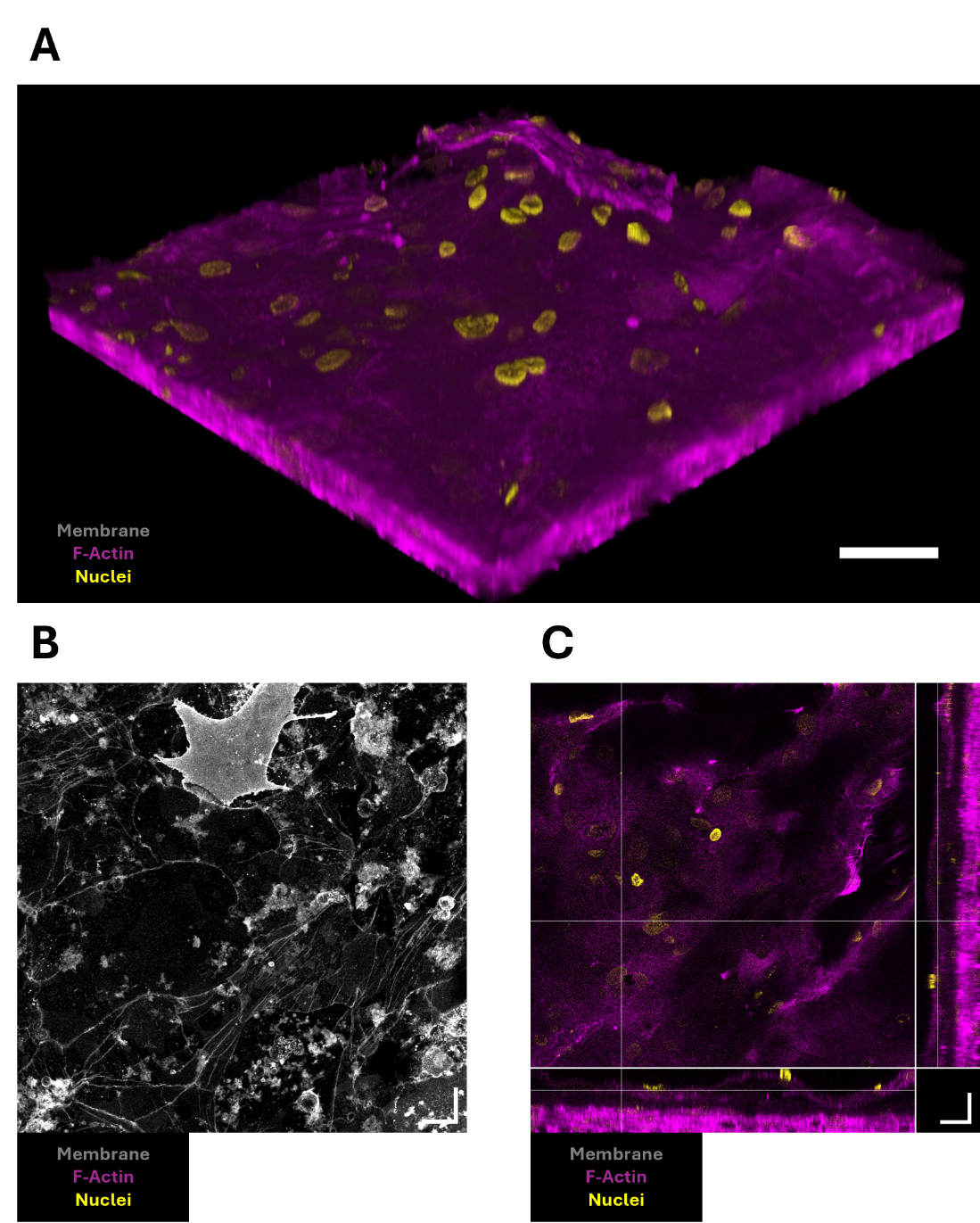

**Supplementary Figure S1:** Confocal imaging of the 3D urine-tolerant human urothelial model in the A**)** top-down (main square) and orthogonal view, **B)** maximum intensity projection and **C)** 3D reconstructed view. Magenta indicates the F-actin or cytoskeleton, yellow indicates cell nuclei and grey indicates the cell membrane of umbrella cells. Scale bar is 20 µm.

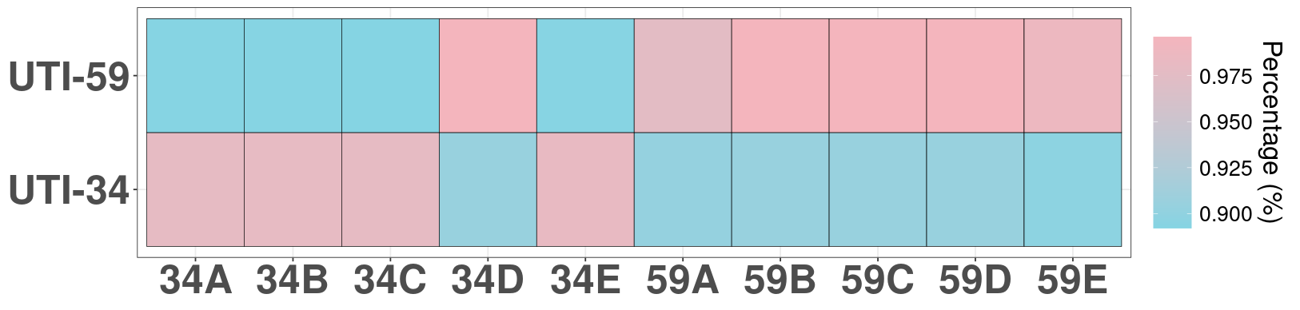

**Supplementary Figure S2:** Average nucleotide identity of the fosfomycin-selected variants compared with their ancestors, UTI-34 and UTI-59.

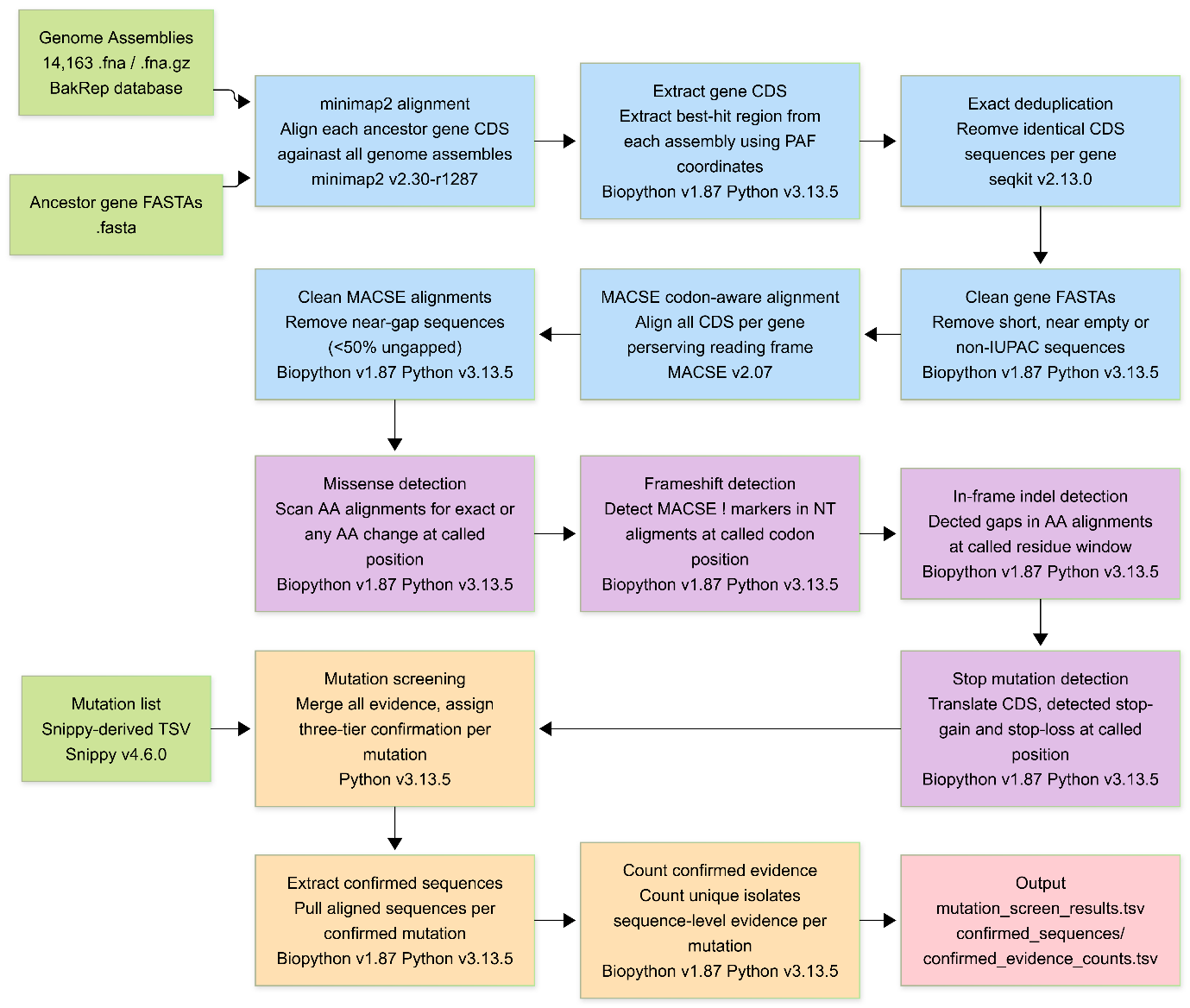

**Supplementary Figure S3:** Workflow for the screening of the mutations observed in the fosfomycin-selected variants against the database of 14,163 *Escherichia coli* genomes isolated from human urine sourced from BakRep. Diagram made using Mermaid.js (<https://mermaid.js.org/>).

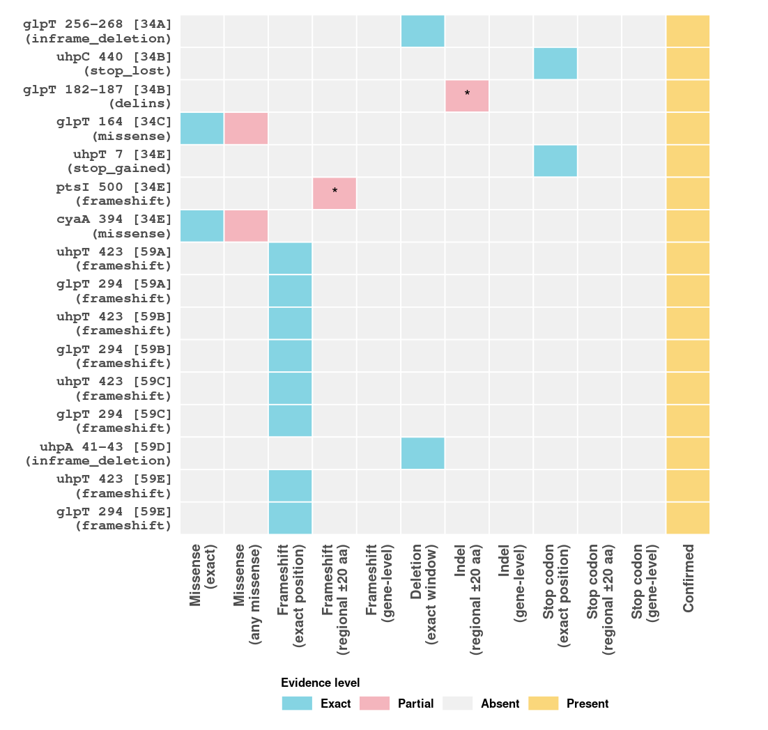

**Supplementary Figure S4:** Presence or absence of the mutations observed in the fosfomycin-selected variants of UTI-34 and UTI-59 against the genomes of the fosfomycin-selected variants. All mutations were correctly called except the deletion-insertion (delins) in *glpT*, where the deletion was called but the cysteine insertion was absent, and *ptsI* frameshift at position 500. Both mutations were confirmed to be present by manual inspection of the aligned sequences with confirmed evidence of the expected mutation, as denoted by a *.

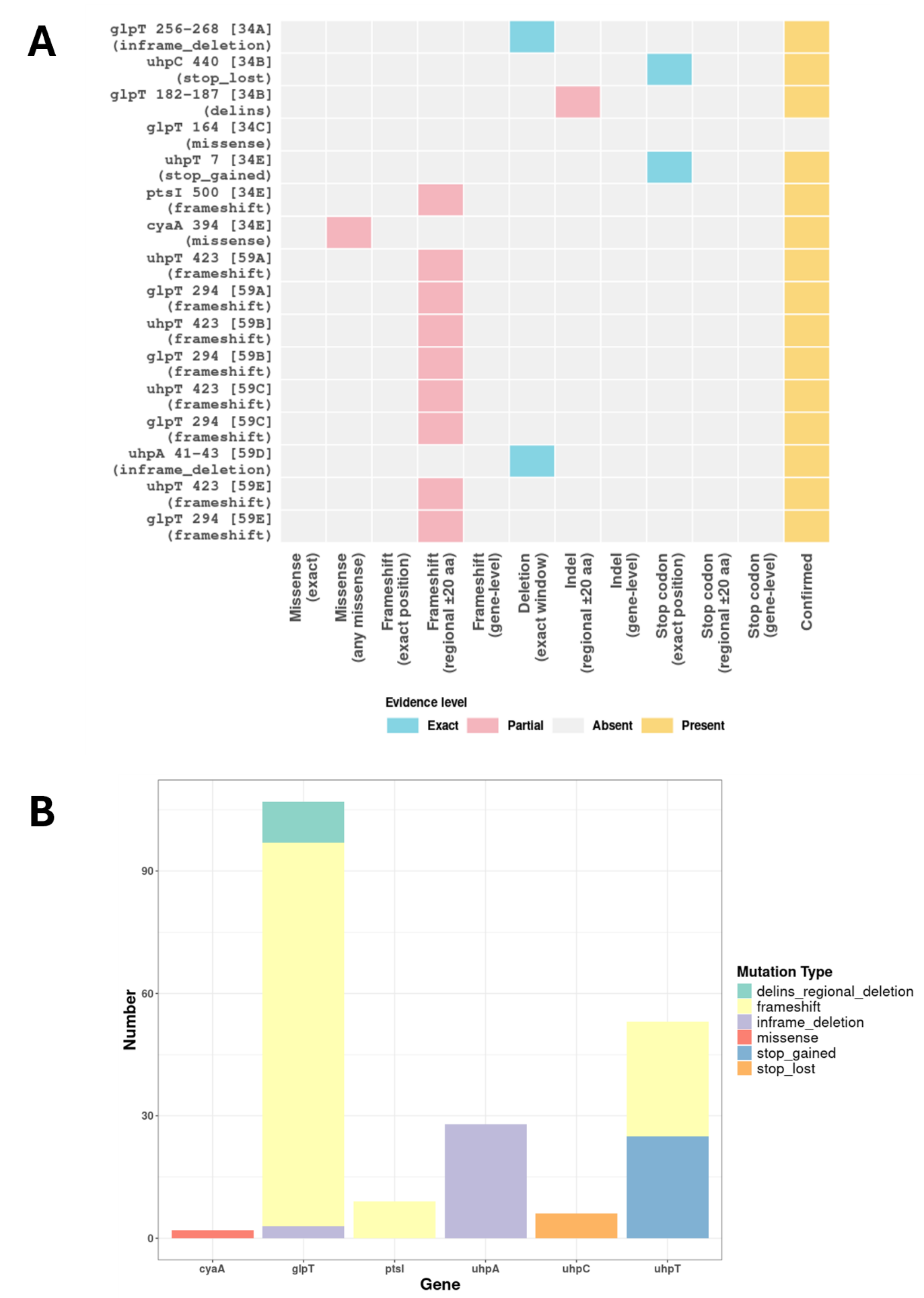

**Supplementary Figure S5:** **A)** Presence or absence of the amino acid changes found in the fosfomycin-selected variants of UTI-34 and UTI-59 in the “all urine” 20,066 *E. coli* genomes isolated from human urine. Exact tier evidence identifies the exact expected mutation found in the fosfomycin-selected variants whereas any missense is a missense at the exact position but a different amino acid change than expected. Regional tier evidence calls the same mutation type 20 amino acids up and downstream of the expected position and gene tier calls the same mutation type anywhere in the gene. **B)** The number of unique genomes with the called tiered mutation within the “all urine” genome dataset per gene.
